# 12/15-Lipoxygenases mediate neuropathic-like pain hypersensitivity in female mice

**DOI:** 10.1101/2024.04.04.588153

**Authors:** B Brown, I Chen, C Miliano, LB Murdaugh, Y Dong, KA Eddinger, TL Yaksh, MD Burton, MW Buczynski, AM Gregus

## Abstract

It is estimated that chronic neuropathic pain conditions exhibit up to 10% prevalence in the general population, with increased incidence in females. However, nonsteroidal inflammatory drugs (NSAIDs) are ineffective, and currently indicated prescription treatments such as opioids, anticonvulsants, and antidepressants provide only limited therapeutic benefit. In the current work, we extended previous studies in male rats utilizing a paradigm of central Toll-like receptor 4 (TLR4)-dependent, NSAID-unresponsive neuropathic-like pain hypersensitivity to male and female C57BL/6N mice, uncovering an unexpected hyperalgesic phenotype in female mice following intrathecal (IT) LPS. In contrast to previous reports in female C57BL/6J mice, female C57BL/6N mice displayed tactile and cold allodynia, grip force deficits, and locomotor hyperactivity in response to IT LPS. Congruent with our previous observations in male rats, systemic inhibition of 12/15-Lipoxygenases (12/15-LOX) in female B6N mice with selective inhibitors – ML355 (targeting 12-LOX-p) and ML351 (targeting 15-LOX-1) – completely reversed allodynia and grip force deficits. We demonstrate here that 12/15-LOX enzymes also are expressed in mouse spinal cord and that 12/15-LOX metabolites produce tactile allodynia when administered spinally (IT) or peripherally (intraplantar in the paw, IPLT) in a hyperalgesic priming model, similar to others observations with the cyclooxygenase (COX) metabolite Prostaglandin E_2_ (PGE_2_). Surprisingly, we did not detect hyperalgesic priming following IT administration of LPS, indicating that this phenomenon likely requires peripheral activation of nociceptors. Collectively, these data suggest that 12/15-LOX enzymes contribute to neuropathic-like pain hypersensitivity in rodents, with potential translatability as druggable targets across sexes and species using multiple reflexive and non-reflexive outcome measures.

## Introduction

Chronic pain syndromes with neuropathic components are extremely challenging to manage as they are refractory to treatment with nonsteroidal inflammatory drugs (NSAIDs), and most patients report inadequate relief from first-line therapies such as anticonvulsants and antidepressants [20]. Long-term use of opioids is not recommended due to potential risk of misuse, diversion and increased overdose mortality rates [4; 68]. Thus, the lack of effective therapeutics for management of chronic neuropathic pain represents a critical unmet need.

It is well-established that the innate and adaptive immune systems contribute to peripheral and central sensitization, and that their relative roles underlying the experience of chronic pain are dependent upon several factors including sex [26; 38]. Infiltrating leukocytes coordinate in the periphery with activated Schwann cells and satellite cells and communicate with CNS resident astrocytes, microglia, and oligodendrocytes, which in turn release mediators including glutamate, substance P, ATP, cytokines and chemokines, and Toll-like receptor (TLR) ligands. These factors sensitize nociceptors and facilitate the transition from acute to chronic pain in the context of disease, metabolic challenge, or injury that cannot be repaired [31; 55]. This acute to chronic transition in neuropathic pain can be modeled by hyperalgesic priming, or a “2-hit” peripheral nociceptive stimulus paradigm [3; 16].

Emerging evidence indicates sex-specific involvement in pain hypersensitivity of TLR4 expressed on particular cell types (macrophages and microglia in males, T-cells and nociceptors in females) [41; 60; 63; 69]. Intrathecal (IT) delivery of lipopolysaccharide (LPS) or endogenous TLR4 ligands (*e.g.*, high mobility group box 1, HMGB1) elicits tactile allodynia that is more pronounced and persists longer in male than female C57BL/6J mice [1; 53; 59; 70]. In contrast, this sex difference in tactile allodynia is absent when LPS is administered via intracerebroventricular or peripheral routes, indicating that this sex-specific effect of LPS occurs only at the spinal level [22; 59].

Previously, we have shown that spinal 12/15-Lipoxygenases (12/15-LOX) are activated during peripheral inflammation [11; 24; 25] and following direct stimulation of TLR4 with IT LPS [23] in male rats. We found that spinal TLR4 activation increases microglial 12/15-LOX metabolites to mediate a neuropathic-like pain state in the absence of peripheral inflammation [23; 54]. Both systemic and IT administration of NSAIDs such as ibuprofen and ketorolac fail to attenuate nociceptive behavior despite complete inhibition of TLR4-induced spinal prostaglandin E_2_ (PGE_2_) release via cyclooxygenases, indicating that 12/15-LOX metabolites contribute directly to NSAID-unresponsive allodynia in rats. However, the role of 12/15-LOX enzymes in neuropathic-like pain states in mice have not been fully examined. There are 6 different 12/15-LOX genes [10]: *Alox12*, *Alox12b*, *Alox12e* (ALOX12P2 in human), *Aloxe3*, *Alox15*, and *Alox15b* (Alox8 in mouse). Herein, we focused on *Alox12* and *Alox15,* the primary enzymes that generate 12(S)-Hydroxyeicosatetraenoic acid (12(S)-HETE) or 15(S)-Hydroxyeicosatetraenoic acid (15(S)-HETE), as these metabolites elicit tactile allodynia directly when administered spinally *in vivo* [24]. In the current report, we examined effects of IT LPS on pain-like behaviors in both sexes in another substrain of BL/6 mice (C57BL/6N), which exhibit differences from C57BL/6J in several behaviors including nociception [9; 67], uncovering an unexpected IT LPS-sensitive pain phenotype in female B6N mice that is completely reversed by selective 12/15-LOX inhibition.

## Methods

### Animals

Male and female C57BL/6N mice (Inotiv; formerly Envigo) or male C57BL/6J mice (Jackson) 12-20 weeks old (20-30g) were used in accordance with protocols approved by the Institutional Animal Care and Use Committees (IACUC) of Virginia Tech, University of Texas at Dallas, and the University of California, San Diego, and all experiments complied with the ARRIVE guidelines. Mice were housed in a minimum of 2 to a maximum of 5 per cage under normal light cycle (7AM-on, 7PM-off) with ad-libitum access to standard chow and water, and all experimental procedures were performed during the light cycle. All behavioral testing was performed during regular light hours 9AM - 5PM by the same observer who was blinded to the treatment groups, with animals randomized by another investigator. The experimental observer was unblinded to treatments at the conclusion of experiments.

### Drug preparation and delivery

#### Drugs and chemicals

Lipopolysaccharide (LPS) from E. coli 0111:B4 strain was purchased from Invivogen (Catalog # tlrl-3pelps). Bioactive lipids prepared in a vehicle (VEH) of sterile, endotoxin-free phosphate buffered saline (PBS, Corning REF#21040-CM) were obtained from Cayman Chemical: Prostaglandin E_2_ (PGE_2_, Catalog # 14010), 12-Hydroxyeicosatetraenoic acid (12(S)-HETE, Catalog #34570), 15-Hydroxyeicosatetraenoic acid (15(S)-HETE, Catalog #34720) and Ketorolac (KETO, Catalog #70690). Inhibitors of 15-LOX-1 (ML351, Cayman Catalog #16119, batch# 0480737-28 or MCE Catalog # HY-111310, lot# 110818) and 12-LOX-p (ML355, Cayman Catalog # 18537, lot# 0555321-8 or MCE Catalog # HY-12341, lot# 29241) were prepared in a 1:1 mix of DMSO (Sigma Catalog # 41640):Tween-20 (Sigma Catalog #P9416) and then diluted in water and PBS for a final dilution of 1:1:1:8, or 5% DMSO/5%Tween-80. No differences were observed in effects of drugs obtained from different companies, so multiple sources are included as an element of rigor for our studies. Isoflurane (NDC 13985-528-60) was obtained from MWI Animal Health.

#### Intrathecal Injections

Following measurement of baseline nociceptive thresholds (see below), percutaneous intrathecal (IT) injections of sterile PBS vehicle (VEH) or LPS were performed as previously described [30; 70]. Briefly, anesthesia (1.5-2% induction; 0.8-1.3% maintenance) was induced in a chamber until loss of the righting reflex was observed using a low-flow vaporizer (Somnosuite, Kent Scientific) or standard vaporizer and then a 30 gauge-1/2” needle attached to a 50 μl Hamilton syringe was inserted percutaneously into the midline between L5 and L6 vertebrae. Proper placement was ensured by observation of a robust tailflick response upon needle insertion. Agents were administered in a volume of 5 μl over an interval of ∼30 seconds. Following recovery from anesthesia, mice were evaluated for normal motor coordination and muscle tone, and any animals exhibiting motor or postural deficits after injection (<1%) were immediately sacrificed. We did not observe any motor dysfunction or loss of muscle tone in mice included in this study, or sustained tactile allodynia in IT saline-treated groups. We assessed multiple nociceptive behavioral output measures including tactile allodynia, cold allodynia, grip force, and open field (see detailed descriptions below) following IT LPS.

#### Intraperitoneal injection

In some experiments, mice received intraperitoneal (IP) injections of either vehicle or 30 mg/kg of ML351 or ML355 (10 ml/kg body weight) as a post-treatment at 5 and 24 h following IT VEH or LPS administration. We assessed multiple nociceptive behavioral output measures including tactile allodynia, cold allodynia, and grip force for up to 48 h following IT LPS.

#### Intraplantar Injections

For hyperalgesic priming, mice were briefly anesthetized with isoflurane via low-flow vaporizer (Somnosuite, Kent Scientific) and then injected with 1 μg LPS/10 μl into the plantar surface of the left hindpaw (day 0). At defined timepoints following IPLT LPS (or IT in some experiments), subthreshold doses (100ng/10uL each) of PGE_2_ (day 14), 15(S)-HETE (day 23) and 12(S)-HETE (day 27) were injected into the left hindpaw. We assessed tactile and cold allodynia as nociceptive behavioral outputs at defined timepoints at baseline, 4h, 1d, and on 14d, 23d, 27d.

#### Plasmids, cell culture, and transfection

cDNAs cloned and inserted into pReceiver-M03 vector driven by CMV promoter with c-terminal GFP were purchased from Genecopoeia (*Alox12*, source accession number NM_007440.5, catalog #EX-Mm01165-M03; *Alox15*, source accession number NM_009660.3, catalog #EX-Mm34152-M03-GS). HEK 293T cells were purchased from ATCC and cultured in high glucose DMEM with 2 mM GlutaMAX and pyruvate (Life Technologies catalog #10567022) supplemented with 10% heat-inactivated characterized fetal bovine serum (Hyclone, Thomas Scientific Catalog #C838T01), and 100 U/ml penicillin G sodium + 100 μg/ml streptomycin Life Technologies Catalog # 15140122). One day prior to transfection, cells were seeded at a density of 8 x 10^6^ cells/well in DMEM without antibiotics on 10 cm plates coated with γ-irradiated poly-D-lysine hydrobromide (Sigma Catalog #P7405). Cells were cotransfected using TransIT 2020 (MirusBio Catalog #5404) and then harvested at 48h post-transfection as described previously [25; 46]. Transfected cell-based overexpression systems were used as positive controls for expression of *Alox12* and *Alox15* in real-time qPCR.

#### Quantitative real-time PCR of mouse 12/15-LOX enzymes

The mouse 12/15-Lipoxygenases encompass a family of six enzymes encoded by six distinct genes: 12-LOX-p (*Alox12*); 12(R)-LOX (*Alox12b*); 12-LOX-e (*Alox12e*, pseudogene); eLOX3 (*Aloxe3*); 15-LOX-1 (*Alox15*); and 8/15-LOX-2 (*Alox15b*) [10]. Of the four 12/15-LOX enzymes expressed in rat spinal cord under basal conditions, we investigated two of these in mouse: *Alox12* (12-LOX-p, which produces 12(S)-HETE and Hepoxilins A3 and B3 in mice and humans) and *Alox15* (15-LOX-1, which produces 12(S)-HETE, 15(S)-HETE, and Hepoxilins A3 and B3 in mice and 15(S)-HETE in humans) [10; 25].

Total RNA was isolated from spinal cord using RNeasy Lipid Tissue Mini Kit (Qiagen Catalog # 74804) and RNase-free DNase kit (Zymo catalog #E1010) and cDNA libraries were generated using iScript cDNA synthesis kit (Biorad catalog #1708890) according to the manufacturer’s instructions. Real-time qPCR with for mouse *Alox12* (12-LOX-p), *Alox15* (15-LOX-1), and *Gapdh* was conducted using prevalidated Taqman primers (Life Technologies catalog #s: *Alox12* Mm00545833_m1 FAM_MGB; *Alox15* Mm00507789_m1 small FAM_MGB; *Gapdh* Mm99999915_g1 VIC_MGB). We confirmed specificity of these primer sets against their respective target genes in HEK-293T overexpression systems as described above and as we reported previously [25]. The comparative C_T_ method was used for relative quantification, and 2^-ΔΔCT^ values were calculated and averaged for each target as described in detail previously [25]. For purposes of clarity, in this paper we refer primarily to the protein nomenclature, with the enzyme family as 12/15-LOX and enzyme activities as 12-lipoxygenase (12-LOX) or 15-lipoxygenase (15-LOX).

### Behavioral Testing

#### Tactile Allodynia

Tactile allodynia was evaluated using manual von Frey filaments with buckling forces between 0.02 and 2 g (Touch Test, Stoelting Co.) applied to the mid-plantar surface of each hindpaw using the up-down method as we have described previously {Chaplan, 1994 #58;Gregus, 2018 #378;Chen, 2023 #469;Murdaugh, 2024 #468}. Any mouse with a basal 50% paw withdrawal threshold (PWT) ≤0.79 g was excluded from the study. For baseline measurements, PWTs from both hindpaws were averaged. Data were expressed as 50% gram thresholds vs time, or as area under the curve (hyperalgesic index % change from baseline).

#### Cold Allodynia

Cold allodynia was evaluated using the fixed temperature model as described with modification [6; 7]. Mice were acclimated to the testing room for 5 days and to the apparatus for 2-3h in a 4-sided acrylic chamber (ePlastics, custom design) with only one transparent wall (3 x 3 x 7.5 inches) until they were calm with paws flat on a 0.25” glass testing surface affixed to the top of a customized IKEA steel stand (Catalog #303.811.83). Thirty minutes before starting the test, mice were removed from the apparatus for injections and the glass plates were cleaned off with a paper towel. Given that both hindpaws are affected by IT and IP injections, cold thresholds were measured using a hand timer by their complete removal from the glass after application of the compressed powdered form of dry ice within a 3 ml syringe (with the tip cut off) placed under the surface below each hindpaw, as previously published [6]. Thresholds for each hindpaw were assessed 3 times with a minimum of 7 minutes resting period for the opposite hindpaw and 15 minutes for the same hindpaw, with a 20 second cutoff time to prevent tissue damage. The highest and lowest threshold was removed, and the remaining four thresholds were averaged together to obtain the mean cold sensitivity response. Data were presented as time-effect curves or as area under the curve (hyperalgesic index % change from baseline), with baseline cold thresholds consistent with the published literature.

#### Grip Force

Changes in grip strength were evaluated using a grip force meter (BIOSEB #BIO-GS3) using BIO-CIS response analysis software according to a previous report {Montilla-Garcia, 2017 #348;Bustin, 2023 #470}. Mice were habituated to the testing room and the metal grid for mice (BIO-GRIPGS, BIOSEB) for 3 days before data collection. Briefly, mice were allowed to grasp a metal grid and their tails were gently pulled by hand for 3 seconds to measure the maximal force exerted by all 4 paws of the mouse before releasing the grid. Each mouse was measured 3 times and then averaged to report mean grip force and delta from baseline. Data are reported as mean change from baseline (prior to IT LPS).

#### Open Field Test

Mice were tested as previously described {Johnson, 2021 #485;Murdaugh, 2024 #468;Chen, 2023 #469}, at 6h post IT saline or LPS. Briefly, they were acclimated to the testing room in their home cage with cage lids open for a minimum of 30 minutes. At the start of testing, mice were placed in the corner of an opaque testing arena (43 cm x 43 cm x 43 cm) under illuminated conditions (200 lux) and their movements were recorded for 10 minutes with an overhead camera. The latency to first enter the center space, amount of time spent in the center space (20 cm x 20 cm) and edge (3.5 cm from each wall) spaces, the total number of entries into the edge and center spaces, and the total distance traveled were measured using ANY-maze tracking software (Stoelting Co., version 5.25).

### Statistical Analysis

Statistical analyses were performed using GraphPad Prism (version 10.1.0) as follows: Two-way repeated measures (timecourse) or standard one-way ANOVA (AUC) for multiple group analysis with Bonferroni *post-hoc* for behavioral data or Dunnett’s *post-hoc* for biochemical data. All data are reported as mean ± SEM, and individual data points are indicated where applicable. The criteria for significance were as follows: *P<0.05, **P<0.01, ***P<0.001, ****P<0.0001.

## Results

### Spinal TLR4 activation produces neuropathic-like tactile allodynia in male and female C57BL/6N mice

We and others have found that C57BL/6J female mice exhibit significantly less spinal TLR4-mediated allodynia than their isogenic male counterparts [59; 61; 70]. Given evidence that B6 substrains exhibit differences in nociception [8; 9; 67], we examined if IT delivery of LPS would produce tactile allodynia in both sexes of B6N mice. As depicted in **Figure 1 A-E**, IT LPS elicits robust tactile allodynia in male and female C57BL/6N mice that is dose-dependent and sustained for at least 24 h. Of note, female C57BL/6N exhibited significantly greater pain hypersensitivity than males at the 0.3 μg dose. To confirm that spinal TLR4 activation is independent of peripheral inflammation as shown previously [23; 54], we demonstrated that pretreatment with an analgesic dose of ketorolac (10 μg IT) that reduces IT LPS-induced spinal prostaglandin release did not prevent the development of allodynia in male C57BL/6J mice (**Figure S1)**, consistent with our previous observations in rats. We did not extend these studies to female C57BL/6J as unlike C57BL/6N mice, females of this strain do not develop significant allodynia in response to IT LPS. Collectively, these results indicate that IT LPS produces a neuropathic-like pain state that is unresponsive to NSAID treatment in both sexes of B6N mice.

**Figure 1.**
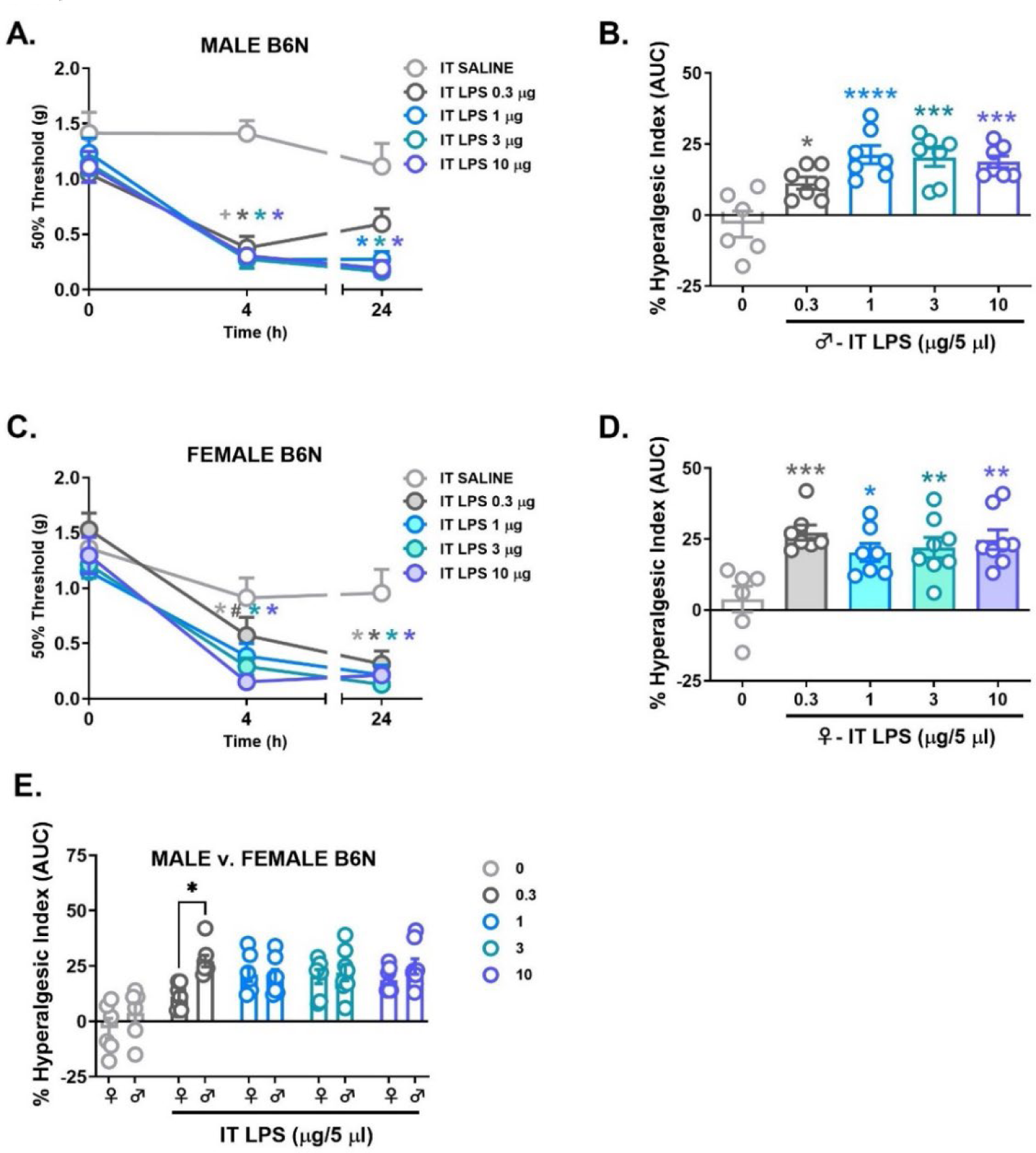
IT LPS produces tactile allodynia in male and female C57BL/6N mice. **(A,C)** Timecourse graphs and **(B,D,E)** AUC values represented as % hyperalgesic index in male or female mice. LPS, Lipopolysaccharide; IT, intrathecal; AUC, Area under the curve. *P<0.05, **P<0.01, ***P<0.001, ****P<0.0001 v. SALINE by Two-way repeated measures ANOVA (timecourse) or One-way ANOVA (AUC) followed by Bonferroni post-hoc test; n = 6-8 mice/group.

### Spinal TLR4 activation produces locomotor hyperactivity in female C57BL/6N mice

We then examined if IT LPS produced effects on anxiety-like behaviors using an affective measure pain hypersensitivity, the open field test. At 6h after IT LPS (when allodynia is maximal), there was a significant effect of sex on some open field behavioral parameters (distance traveled p<0.0001, time immobile p<0.0001, number of center entries p<0.05, and number of edge entries p<0.05), wherein females generally spent less time immobile, traveled greater distance and exhibited a higher number of center and edge entries than males (**Table 1**). However, there was no significant effect of sex, IT LPS dose, or their interaction on center time, edge time, latency to enter the center or edges of the open field, or number of fecal boli, indicating a lack of anxiety-like behavior in this test. Interestingly, despite exhibiting greater allodynia at the 0.3 μg dose of IT LPS, immobility time in females was significantly lower than in males, consistent with increased locomotor activity as measured by distance traveled. These findings indicate that while spinal delivery of LPS at doses up to 10 μg does not produce anxiety-like behaviors in open field for C57BL/6N mice, females exhibit locomotor hyperactivity compared with males at a low dose of endotoxin.

**Table 1.** IT LPS produces sex-specific effects on open field behavior in male and female C57BL/6N mice. At each dose of IT LPS (0, 0.3, 1, 3, and 10 μg), the following parameters were recorded: Distance traveled (m), Time immobile (s), Center time (s), Edge time (s), Number of center entries, Number of edge entries, Latency to center entries (s), Latency to edge entries (s), and Number of fecal boli. IT, intrathecal; LPS, Lipopolysaccharide. *P<0.05, ***P<0.001, ****P<0.0001 males versus females by Two-way ANOVA followed by Bonferroni post-hoc test; n = 7-8 mice/group.

| IT LPS Dose (μg) | 0 | 0.3 | 1 | 3 | 10 | Main effect? |
| --- | --- | --- | --- | --- | --- | --- |
| Distance Traveled (m) | 37.19 ± 2.72 | 32.25 ± 3.37 | 37.46 ± 3.31 | 29.49 ± 2.44 | 32.95 ± 2.67 | Sex $p < 0.0001$; |
| | 43.57 ± 2.71 | 47.30 ± 3.61 | 44.59 ± 2.39 | 43.99 ± 4.71 | 41.26 ± 2.20 | Dose ns; Interaction ns |
| Time Immobile (s) | 74.00 ± 13.21 | 105.51 ± 24.27 | 98.27 ± 17.17 | 131.93 ± 21.15 | 115.06 ± 17.94 | Sex $p < 0.0001$; |
| | 67.19 ± 8.60 | 48.10 ± 6.70 | 59.60 ± 5.90 | 63.95 ± 11.54 | 67.14 ± 11.56 | Dose ns; Interaction ns |
| Center Time (s) | 62.97 ± 13.43 | 72.07 ± 13.78 | 38.43 ± 7.18 | 64.66 ± 9.99 | 68.31 ± 13.80 | Sex, Dose, Interaction n.s. |
| | 58.53 ± 11.83 | 74.61 ± 7.09 | 64.30 ± 13.30 | 65.83 ± 4.71 | 66.06 ± 12.39 | |
| Edge Time (s) | 491.43 ± 24.76 | 493.21 ± 19.85 | 526.83 ± 14.68 | 496.63 ± 21.74 | 478.89 ± 23.08 | Sex, Dose, Interaction n.s. |
| | 495.50 ± 15.27 | 471.19 ± 7.32 | 488.26 ± 14.30 | 478.84 ± 7.78 | 482.40 ± 15.89 | |
| Center Entries (#) | 25.57 ± 3.73 | 24.71 ± 4.78 | 21.29 ± 2.91 | 23.00 ± 4.12 | 26.00 ± 4.09 | Sex $p < 0.05$; |
| | 29.13 ± 3.70 | 34.25 ± 3.74 | 27.38 ± 4.28 | 31.88 ± 4.49 | 28.13 ± 3.29 | Dose ns; Interaction ns |
| Edge Entries (#) | 26.29 ± 3.73 | 25.57 ± 4.74 | 21.43 ± 2.78 | 23.71 ± 4.11 | 26.71 ± 3.93 | Sex $p < 0.05$; |
| | 29.38 ± 3.63 | 35.13 ± 3.78 | 28.00 ± 4.22 | 32.63 ± 4.46 | 28.63 ± 3.13 | Dose ns; Interaction ns |
| Center Entries (latency) | 38.83 ± 8.87 | 43.79 ± 11.74 | 47.21 ± 20.35 | 39.83 ± 17.00 | 35.83 ± 8.78 | Sex, Dose, Interaction n.s. |
| | 23.24 ± 6.06 | 19.46 ± 2.18 | 51.15 ± 16.24 | 47.68 ± 8.68 | 34.46 ± 8.77 | |
| Edge Entries (latency) | 9.83 ± 3.56 | 12.53 ± 2.39 | 11.91 ± 5.85 | 9.17 ± 3.07 | 12.30 ± 4.08 | Sex, Dose, Interaction n.s. |
| | 15.58 ± 7.29 | 7.80 ± 1.57 | 5.80 ± 1.44 | 7.41 ± 1.72 | 10.70 ± 1.47 | |
| Fecal Boli (#) | 0.29 ± 0.29 | 2.71 ± 0.84 | 2.00 ± 0.85 | 1.43 ± 1.02 | 1.57 ± 0.57 | Sex, Dose, Interaction n.s. |
| | 1.00 ± 0.63 | 1.38 ± 0.46 | 2.38 ± 0.56 | 1.75 ± 0.41 | 1.38 ± 0.82 | |

M v F: 0.3 μg $p < 0.001$

### Spinal 12/15-LOX activation contributes to pain hypersensitivity in female C57BL/6N mice

We have shown previously that in male rats, spinal delivery of LPS increases 12/15-lipoxygenase metabolites [23] that directly elicit nociception [24], yet their effects have not yet been examined in mice. As most chronic pain conditions occur more commonly in women [26], we focused on female mice for the remainder of this study. We administered 12(S)-HETE or 15(S)-HETE at 2 different doses (100 ng and 1 μg IT) compared with saline vehicle and then measured their effects on tactile thresholds in female C57BL/6N mice. We observed that IT 12(S)-HETE or 15(S)-HETE produced a dose-dependent, robust tactile allodynia that was sustained for up to 3 days after a single injection at the high dose (**Figure 2 A,B**). Next, we asked if the 12/15-LOX enzymes primarily responsible for synthesizing 12(S)-HETE (*Alox12,* encoding 12-LOX-p) and 15(S)-HETE (*Alox15*, encoding 15-LOX-1) are basally expressed at the spinal level in mice. Quantitative RT-PCR analysis revealed that both *Alox12* and *Alox15* are expressed, albeit at low basal levels in spinal cord of female B6N mice (**Figure 3 A-C**).

**Figure 2.**
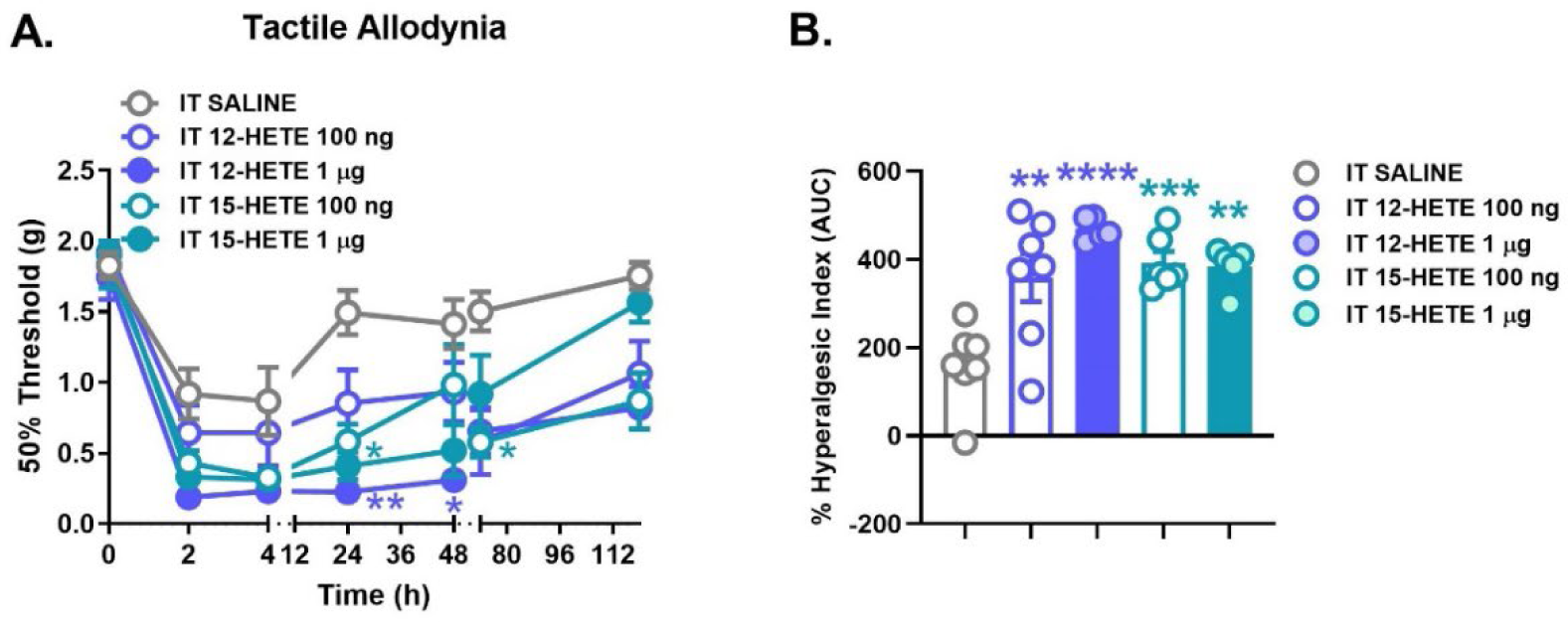
IT delivery of 12/15-LOX metabolites produces robust, persistent tactile allodynia in female C57BL/6N mice. **(A)** Timecourse graph and **(B)** AUC values represented as % hyperalgesic index in female mice. IT, intrathecal; AUC, Area under the curve. 12-Hydroxyeicosatetraenoic acid (12(S)-HETE), 15-Hydroxyeicosatetraenoic acid (15(S)-HETE). *P<0.05, **P<0.01, ***P<0.001, ****P<0.0001 v. SALINE by Two-way repeated measures ANOVA (timecourse) or One-way ANOVA (AUC) followed by Bonferroni post-hoc test; n = 5-7 mice/group.

**Figure 3.**
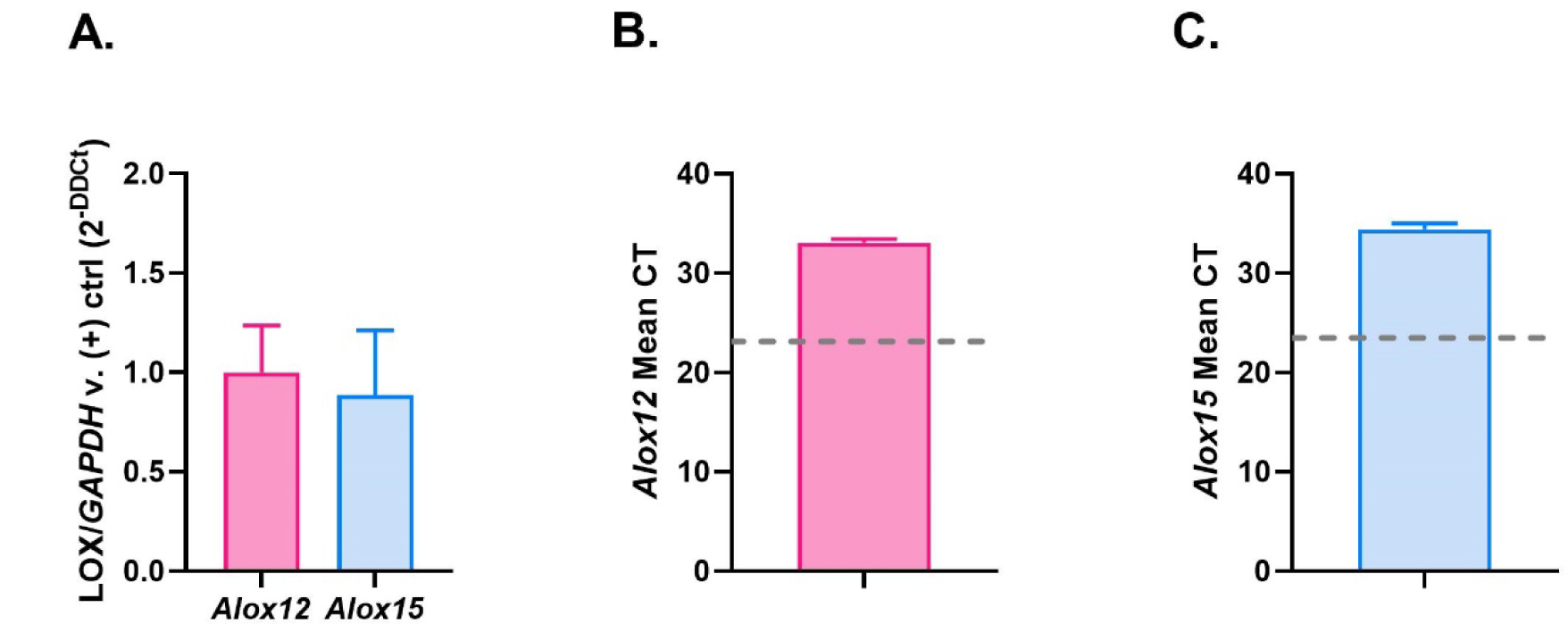
Basal expression of 12/15-LOX enzymes in spinal cord of female C57BL/6N mice. **(A)** Expression of *Alox12* and *Alox15* in mouse spinal cord normalized to GAPDH loading control and positive control cell lines (HEK293T cells overexpressing mouse *Alox12* or *Alox15*) as measured by qPCR. Mean CT values of **(B)** *Alox12* and **(C)** *Alox15* in spinal cord, with mean CT values of respective positive controls marked by gray dotted line. n = 6 mice/group.

### Spinal TLR4-mediated neuropathic-like pain states are mediated by 12/15-LOX activation in female C57BL/6N mice

We have demonstrated previously that pretreatment with inhibitors of 12-LOX-p (ML127) or 15-LOX-1 (ML351) attenuated IT LPS-induced tactile allodynia in a dose-dependent manner in male rats [23]. However, the efficacy of 12/15-LOX inhibitors in reversing established neuropathic-like pain states in mice has not yet been examined. Thus, we investigated if systemic inhibition of 12-LOX-p (with the newer generation molecule ML355) or of 15-LOX-1 (with ML351) could reverse established pain hypersensitivity (**A,B** tactile allodynia; **C,D** cold allodynia; **E,F** grip force deficits) following acute administration of 1 μg IT LPS (**Figure 4**). Timecourse plots revealed that IT LPS-induced pain-like behaviors were maximal at 24h post-injection for all 3 output measures (**A,** tactile; **C,** cold; **E,** grip). Importantly, post-treatment with either ML355 or ML351 (30 mg/kg i.p.) reversed IT LPS-induced tactile allodynia at 6 and 24h (**A,B**) as well as cold allodynia (**C-D**) and grip force deficits at 24h (**E-F**).

**Figure 4.**
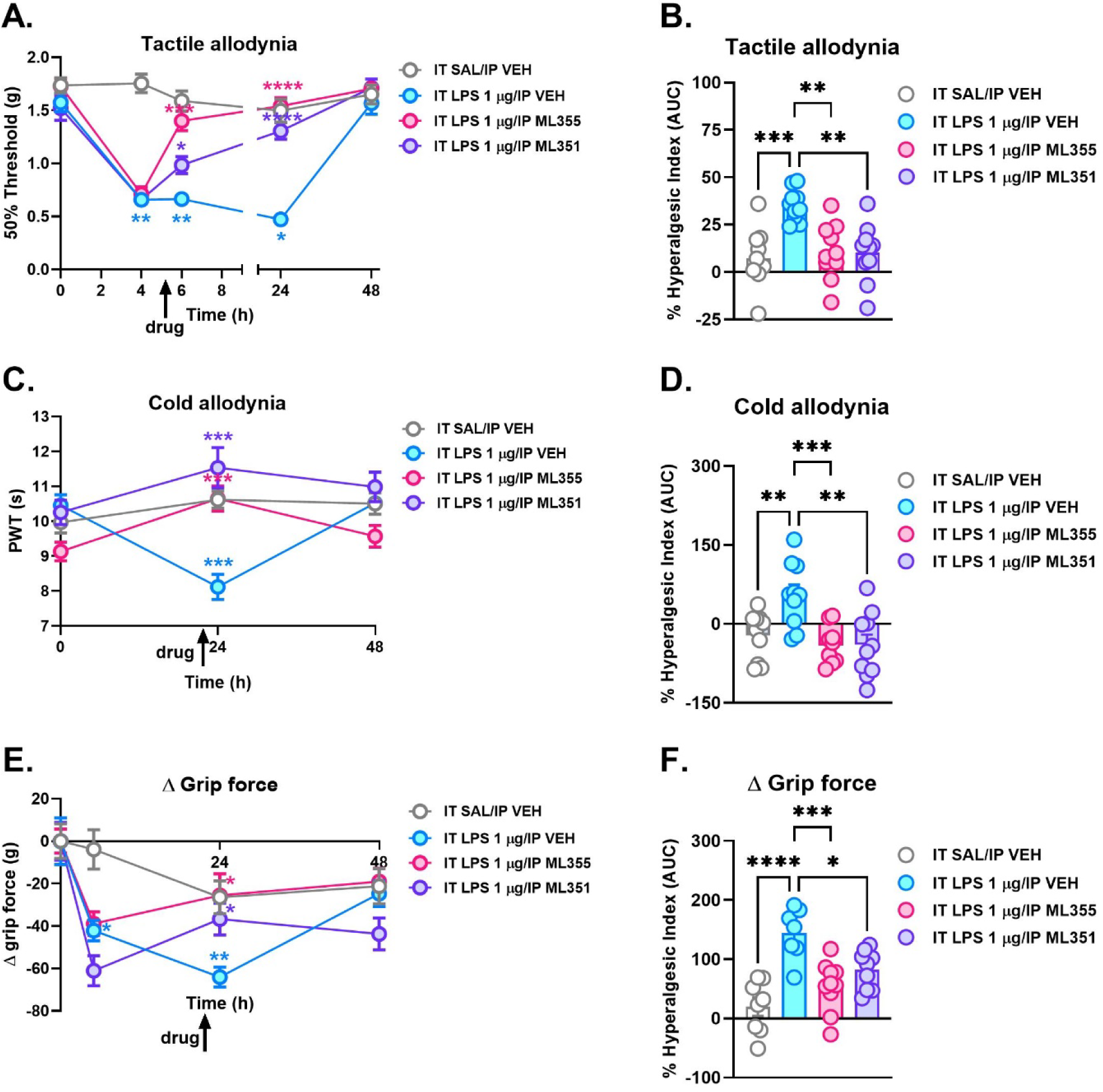
Inhibition of 12- and 15-LOX enzymes reverses IT LPS-induced allodynia and grip force deficits in female C57BL/6N mice. **(A,C,E)** Timecourse and **(B,D,F)** % hyperalgesic index or AUC values for 1 ug IT LPS-induced tactile allodynia **(A,B)**, cold allodynia **(C,D)** and **(E,F)** grip force deficits. IT, intrathecal; LPS, Lipopolysaccharide; SAL, Saline; VEH, Vehicle - 1:1:1:8 DMSO, Tween-80, Saline, Water; AUC, Area under the curve. *P<0.05, **P<0.01, ***P<0.001, ****P<0.0001 v. IT SAL/VEH or IT LPS/VEH by Two-way repeated measures ANOVA (timecourse) or One-way ANOVA (AUC) followed by Bonferroni post-hoc test; n = 10 mice/group.

Next, we examined if 12/15-LOX metabolites contribute to hyperalgesic priming, a model of the acute to chronic transition of pain in rodents [51]. Consistent with previous observations in C57BL/6J mice, IPLT delivery of a subthreshold dose of PGE_2_ (100 ng) at 2-4 weeks after IPLT LPS (1 μg) produced profound tactile [64] but not cold allodynia (**Figure 5 A-C**). Likewise, subthreshold dosing (100 ng) of 15(S)-HETE (**Figure 5 D-F**) or 12(S)-HETE (**Figure 5 G-I**) after paw LPS also evoked sustained tactile hypersensitivity for at least 24 hours after a single IPLT injection. It is important to note we found that IPLT administration of 100 ng of PGE_2_, 12(S)-HETE, or 15(S)-HETE at 14 days after IPLT SAL did not alone elicit tactile or cold allodynia in female C57BL/6N mice in the absence of a priming stimulus, confirming this is a subthreshold dose when administered in the periphery **(Figure S2)**. Finally, we interrogated if hyperalgesic priming could be initiated in the central nervous system (CNS) by IT administration of LPS, followed 2 weeks later by a subthreshold dose of either IT (central) or IPLT (peripheral) PGE_2_ (100 ng). Surprisingly, we found that neither route of administration of PGE_2_ elicited a behavioral effect when priming was initiated by a central delivery of LPS **(Figure S3)**. Collectively, these results demonstrate that 12/15-LOX enzymes contribute to the development and maintenance of spinal TLR4-dependent, NSAID-unresponsive pain hypersensitivity in female mice.

**Figure 5.**
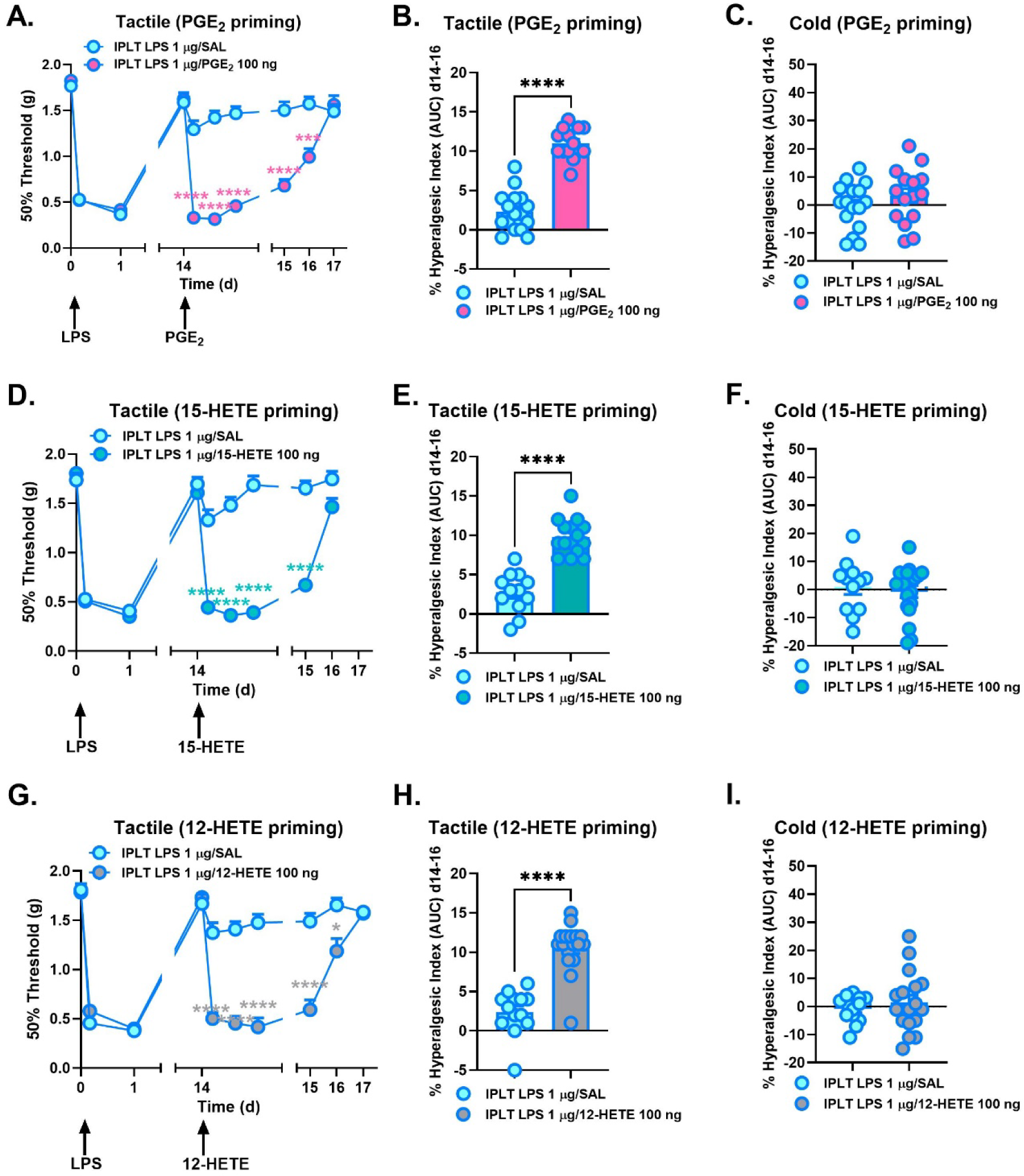
Hyperalgesic priming with intraplantar LPS and either Cyclooxygenase or 12/15-Lipoxygenase metabolites produces tactile but not cold allodynia. Hyperalgesic priming by IPLT delivery of **(A-C)** PGE_2_, **(D-F)** 15(S)-HETE, **(G-I)** or 12(S)-HETE (100 ng) 2-4 weeks after IPLT LPS (1 μg). **(A,D,G)** Timecourse or **(B,C,E,F, H,I)** % hyperalgesic index or AUC values. IPLT, intraplantar; LPS, Lipopolysaccharide; SAL, Saline; AUC, Area under the curve. *P<0.05, ***P<0.001, ****P<0.0001 v. IPLT LPS/SAL by Two-way repeated measures ANOVA (timecourse) or One-way ANOVA (AUC) followed by Bonferroni post-hoc test; n = 12-18 mice/group.

## Discussion

In this study, we extended our previous work utilizing a paradigm of a spinally-mediated TLR4-dependent and NSAID-unresponsive neuropathic-like pain state, uncovering an unexpectedly robust, persistent pain hypersensitivity in female C57BL/6N mice [23; 54]. Our findings in female C57BL/6N mice contrast with previous observations in female mice of a different B6 substrain (C57BL/6J), which do not develop significant allodynia following IT LPS [1; 53; 59; 70]. Other groups also have described significant differences in measures of nociception between these substrains. For example, male B6J mice exhibit lower baseline thermal withdrawal latencies than male B6N mice [9; 42; 58]. However, mixed sex groups of B6N mice develop more pronounced formalin-induced tactile allodynia and anxiety-like behaviors than mixed sex groups of B6J mice [67], suggesting potentially greater sensitivity of female B6N mice to at least some modalities of injury-induced noxious stimuli. In addition, we have shown previously that the delayed tactile allodynia following IPLT formalin is TLR4-dependent in male and female B6 mice from Harlan [70], so our findings regarding the development of IT LPS-induced allodynia, grip deficits, and locomotor hyperactivity are congruent with the literature. While the molecular mechanisms for these substrain differences in nociception are as yet unclear, there are a number of differentially expressed candidate genes between B6J and B6N lines that could be explored in future studies including, but not limited to: (1) nicotinamide nucleotide transhydrogenase (*Nnt*), an enzyme that couples hydride transfer between NAD(H) and NADP(+) to proton translocation across the inner mitochondrial membrane [21; 42; 47; 58]; (2) WD repeat and FYVE domain containing 1 (*Wdfy1*), which mediates TLR3/TLR4 signaling through TRIF [29; 47]; (3) dynein light chain Tctex-type 1 (*Dynlt1*), which regulates mitochondrial permeabilization in hypoxia [17; 47]; and (4) RAB4A, member RAS oncogene family (*Rab4A*), a small GTPase associated with early endosomes which regulates membrane trafficking and recycling of several cell-surface proteins such as glucagon receptors, neurokinin 1 (NK1) receptors, and GLUT4 glucose transporters [34; 35; 47; 52].

In addition to evoked testing of pain-like behaviors (tactile and cold allodynia), we also examined a functional output (grip force deficit) and non-reflexive outcome measure (open field activity). Indeed, a major criticism of preclinical models (and a potential source of failure for investigational new drugs in Phase II) is the principal focus on evoked measures of pain in which the primary measure is an escape response or avoidance paradigm, which bears little relevance to the clinical situation of ongoing spontaneous pain [72]. Thus, the inclusion of non-reflexive outcome measures in preclinical target ID and validation studies is crucial. As expected, IT LPS produced significant deficits in grip force [12; 36] that were reversed by 12/15-LOX inhibition. While IT LPS elicited sex-specific locomotor hyperactivity in female mice, there were minimal effects on anxiety-like behaviors in this test. Others have observed mixed effects on anxiety-like behaviors following LPS, with some detecting an increase and others demonstrating no change [5; 15; 27; 28; 37; 43; 56; 62]. These studies examined behavioral effects of systemic (but not spinal) administration of LPS at different doses and timepoints post-injection in different ages, all of which are factors that may account for discrepant LPS-mediated effects on anxiety phenotypes in rodents. While CSF is cycled to supraspinal sites after IT injection [50], it is possible that this route of administration does not result in high enough concentrations of LPS in relevant brain areas to affect anxiety. Alternatively, it is possible that although the timepoint we examined (6h post-injection) is one of maximal allodynia expression, it may not be optimal for detection of anxiety-like behaviors. However, this scenario is less likely as others observed systemic LPS-induced anxiety-like behaviors anywhere from 2h to 1 month post-injection. Had we observed an anxious phenotype following IT LPS, we would expect 12/15-LOX inhibition to attenuate this effect, as others have observed that overexpression of 12/15-LOX increases anxiety-like behaviors in female mice [33].

Most importantly, the present work demonstrates that 12/15-LOX enzymes contribute to neuropathic pain-like hypersensitivity in female mice, which corroborates and extends our previous observations in male rats [23], supporting potential translatability of this mechanism across species and sexes. Drugs targeting pathways engaged in multiple species during a disease state exhibit higher probability for successful translation into human treatments [48; 71]. Furthermore, it is vital to include females in preclinical studies of pain as chronic pain conditions (particularly neuropathies with autoimmune origin) occur more commonly in women [19; 26; 44]. Our observations also complement accumulating evidence in support of a critical role for 12/15-LOX activation in hyperalgesia observed in preclinical studies from several groups [2; 11; 39; 40; 57]. Herein, we show that 12/15-LOX metabolites 12(S)-HETE and 15(S)-HETE elicit tactile allodynia in the periphery (after IPLT administration) and in the CNS (after IT administration) of female mice. Additionally, 12/15-LOX metabolites produce tactile but not cold allodynia when administered at subthreshold doses in the periphery in a hyperalgesic priming paradigm initiated with IPLT LPS. These results corroborate previous observations of sustained tactile allodynia following hyperalgesic priming with PGE_2_ [18; 49; 51; 64], which we replicate in the current study, and minimal (if any) effect of COX or 12/15-LOX metabolites on thermal thresholds [2; 24; 65; 66]. We have shown previously in male rats that in addition to COX metabolites such as PGE_2_, IT LPS triggers increased production of 12/15-LOX metabolites in spinal microglia and astrocytes, and that the concurrent tactile allodynia is not alleviated by analgesic doses of NSAIDs [23; 54]. In contrast, inhibition of 12-LOX-p or 15-LOX-1 prevents [23] and reverses (current study) this pain hypersensitivity (tactile and cold allodynia, grip deficits) in rodents. The efficacy of ML351 and ML355 in reversing cold allodynia is unexpected given the observed lack of effect of individual 12/15-LOX metabolites on cold thresholds. It is possible that cold allodynia requires action of multiple 12/15-LOX metabolites, elements of signaling pathways engaged downstream of 12/15-LOX activation following IT LPS, or that altered cold thresholds represent off-target effects of the inhibitors at higher doses – mechanisms that could be interrogated in future studies. Taken together, these results support 12/15-LOX enzymes as potentially viable druggable targets for neuropathic pain states and thereby warrant further study.

## Acknowledgements

This work was supported by NIH grants NIAMS R01 AR075241 (AMG); NIDA R00 DA035865 and NCI R01 CA284075 (MWB); NIGMS R35 GM147094 and NIDDK R21 DK130015 (MDB); NINDS R01 NS099338 and NINDS R01 NS132483 (TLY)

## COI statement

The authors have no competing financial interests to declare.

---

**Figure S1.**
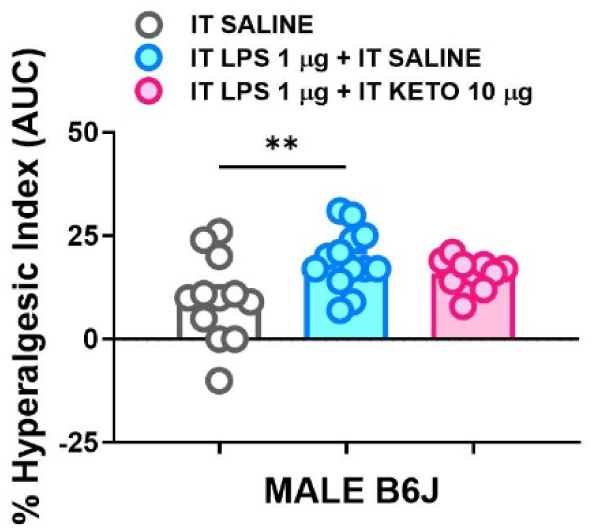
IT LPS produces tactile allodynia that is unresponsive to NSAIDs. AUC values represented as % hyperalgesic index in male mice. LPS, Lipopolysaccharide; IT, intrathecal; KETO, ketorolac; AUC, Area under the curve. **P<0.01 v. SALINE by One-way ANOVA (AUC) followed by Bonferroni post-hoc test; n = 9-13 male mice/group.

**Figure S2.**
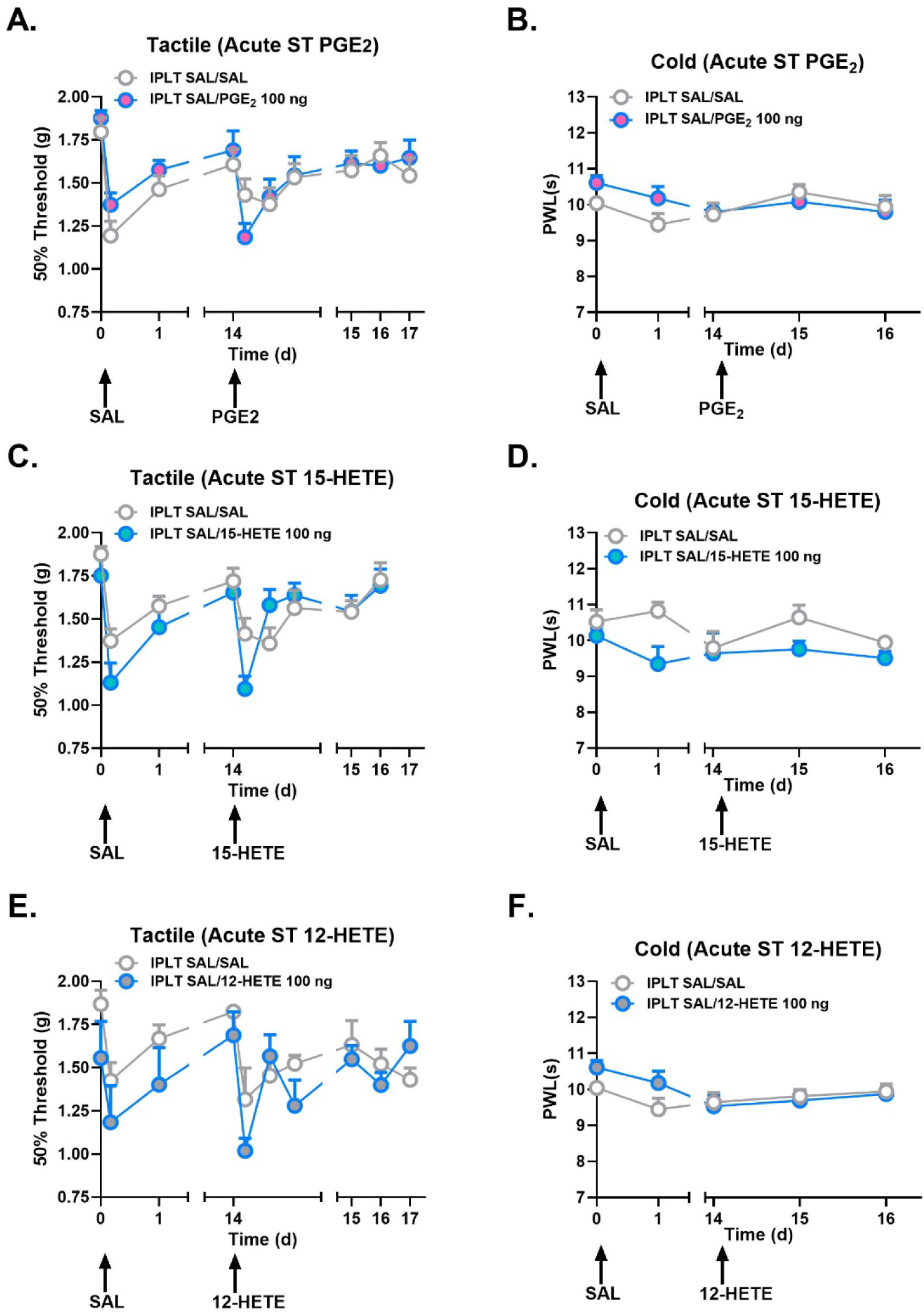
Subthreshold doses of Cyclooxygenase and 12/15-Lipoxygenase metabolites delivered peripherally do not produce sustained tactile or cold allodynia in the absence of priming. Timecourses for IPLT delivery of **(A-C)** PGE_2_, **(D-F)** 15(S)-HETE, **(G-I)** or 12(S)-HETE (100 ng) 2-4 weeks after IPLT SAL. IPLT, intraplantar; SAL, Saline. P values calculated by Two-way repeated measures ANOVA followed by Bonferroni post-hoc test; n = 10 mice/group.

**Figure S3.**
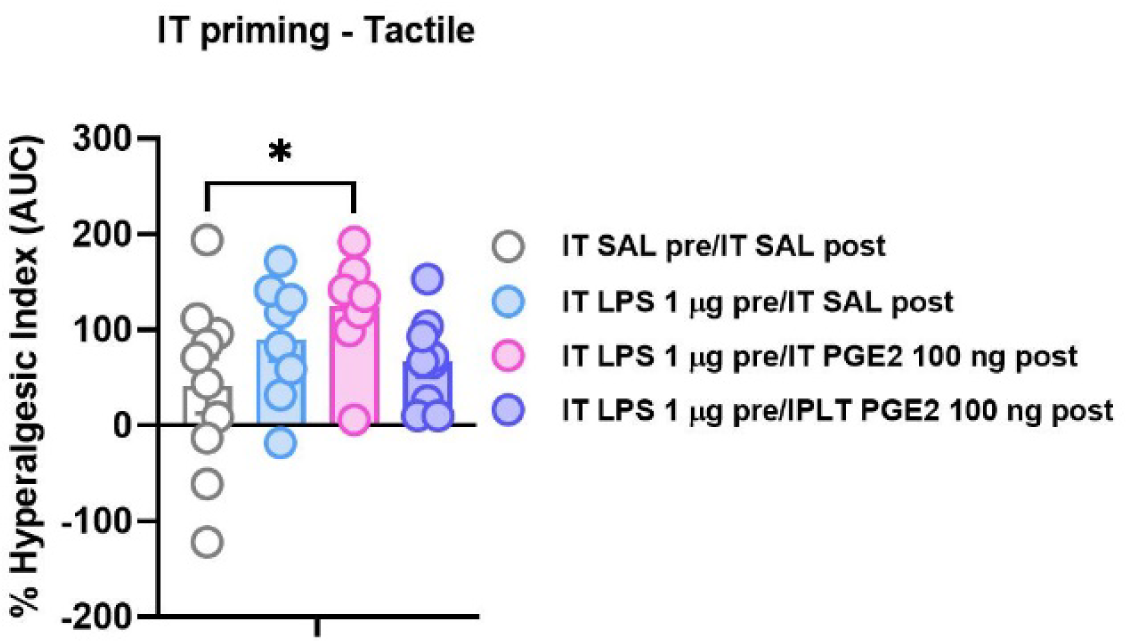
Lack of hyperalgesic priming with intrathecal LPS and either intrathecal or intraplantar PGE_2_. AUC for IT delivery of SAL or LPS followed 2 weeks later by IT SAL, IT PGE_2_ or IPLT PGE_2_. IPLT, intraplantar; IT, intrathecal; SAL, Saline. P values calculated by Two-way repeated measures ANOVA followed by Bonferroni post-hoc test; n = 8-10 mice/group.

## Notes

### Competing Interest Statement

The authors have declared no competing interest.

## References.

1. [1] Agalave NM, Rudjito R, Farinotti AB, Khoonsari PE, Sandor K, Nomura Y, Szabo-Pardi TA, Urbina CM, Palada V, Price TJ, Erlandsson Harris H, Burton MD, Kultima K, Svensson CI. Sex-dependent role of microglia in disulfide high mobility group box 1 protein-mediated mechanical hypersensitivity. Pain 2021;162(2):446–458.

2. [2] Aley O, Levine JD. Contribution of 5- and 12-lipoxygenase products to mechanical hyperalgesia induced by prostaglandin E(2) and epinephrine in the rat. Exp Brain Res 2003;148(4):482–487.

3. [3] Araldi D, Bogen O, Green PG, Levine JD. Role of Nociceptor Toll-like Receptor 4 (TLR4) in Opioid-Induced Hyperalgesia and Hyperalgesic Priming. J Neurosci 2019;39(33):6414–6424.

4. [4] Balanaser M, Carley M, Baron R, Finnerup NB, Moore RA, Rowbotham MC, Chaparro LE, Gilron I. Combination pharmacotherapy for the treatment of neuropathic pain in adults: systematic review and meta-analysis. Pain 2023;164(2):230–251.

5. [5] Bassi GS, Kanashiro A, Santin FM, de Souza GE, Nobre MJ, Coimbra NC. Lipopolysaccharide-induced sickness behaviour evaluated in different models of anxiety and innate fear in rats. Basic Clin Pharmacol Toxicol 2012;110(4):359–369.

6. [6] Brenner DS, Golden JP, Gereau RWt. A novel behavioral assay for measuring cold sensation in mice. PLoS One 2012;7(6):e39765.

7. [7] Brenner DS, Golden JP, Vogt SK, Gereau RWt. A simple and inexpensive method for determining cold sensitivity and adaptation in mice. J Vis Exp 2015(97).

8. [8] Bryant CD, Bagdas D, Goldberg LR, Khalefa T, Reed ER, Kirkpatrick SL, Kelliher JC, Chen MM, Johnson WE, Mulligan MK, Imad Damaj M. C57BL/6 substrain differences in inflammatory and neuropathic nociception and genetic mapping of a major quantitative trait locus underlying acute thermal nociception. Mol Pain 2019;15:1744806918825046.

9. [9] Bryant CD, Zhang NN, Sokoloff G, Fanselow MS, Ennes HS, Palmer AA, McRoberts JA. Behavioral differences among C57BL/6 substrains: implications for transgenic and knockout studies. J Neurogenet 2008;22(4):315–331.

10. [10] Buczynski MW, Dumlao DS, Dennis EA. Thematic Review Series: Proteomics. An integrated omics analysis of eicosanoid biology. J Lipid Res 2009;50(6):1015–1038.

11. [11] Buczynski MW, Svensson CI, Dumlao DS, Fitzsimmons BL, Shim JH, Scherbart TJ, Jacobsen FE, Hua XY, Yaksh TL, Dennis EA. Inflammatory hyperalgesia induces essential bioactive lipid production in the spinal cord. J Neurochem 2010;114(4):981–993.

12. [12] Bustin KA, Shishikura K, Chen I, Lin Z, McKnight N, Chang Y, Wang X, Li JJ, Arellano E, Pei L, Morton PD, Gregus AM, Buczynski MW, Matthews ML. Phenelzine-based probes reveal Secernin-3 is involved in thermal nociception. Mol Cell Neurosci 2023;125:103842.

13. [13] Chaplan SR, Bach FW, Pogrel JW, Chung JM, Yaksh TL. Quantitative assessment of tactile allodynia in the rat paw. J Neurosci Methods 1994;53(1):55–63.

14. [14] Chen I, Murdaugh LB, Miliano C, Dong Y, Gregus AM, Buczynski MW. NAPE-PLD regulates specific baseline affective behaviors but is dispensable for inflammatory hyperalgesia. Neurobiol Pain 2023;14:100135.

15. [15] Clark SM, Michael KC, Klaus J, Mert A, Romano-Verthelyi A, Sand J, Tonelli LH. Dissociation between sickness behavior and emotionality during lipopolysaccharide challenge in lymphocyte deficient Rag2(-/-) mice. Behav Brain Res 2015;278:74–82.

16. [16] Cunha TM, Verri WA, Jr., Fukada SY, Guerrero AT, Santodomingo-Garzon T, Poole S, Parada CA, Ferreira SH, Cunha FQ. TNF-alpha and IL-1beta mediate inflammatory hypernociception in mice triggered by B1 but not B2 kinin receptor. Eur J Pharmacol 2007;573(1-3):221–229.

17. [17] Fang YD, Xu X, Dang YM, Zhang YM, Zhang JP, Hu JY, Zhang Q, Dai X, Teng M, Zhang DX, Huang YS. MAP4 mechanism that stabilizes mitochondrial permeability transition in hypoxia: microtubule enhancement and DYNLT1 interaction with VDAC1. PLoS One 2011;6(12):e28052.

18. [18] Ferrari LF, Levine JD. Plasma membrane mechanisms in a preclinical rat model of chronic pain. J Pain 2015;16(1):60–66.

19. [19] Fillingim RB, King CD, Ribeiro-Dasilva MC, Rahim-Williams B, Riley JL, 3rd. Sex, gender, and pain: a review of recent clinical and experimental findings. J Pain 2009;10(5):447–485.

20. [20] Finnerup NB, Attal N, Haroutounian S, McNicol E, Baron R, Dworkin RH, Gilron I, Haanpaa M, Hansson P, Jensen TS, Kamerman PR, Lund K, Moore A, Raja SN, Rice AS, Rowbotham M, Sena E, Siddall P, Smith BH, Wallace M. Pharmacotherapy for neuropathic pain in adults: a systematic review and meta-analysis. Lancet Neurol 2015;14(2):162–173.

21. [21] Fontaine DA, Davis DB. Attention to Background Strain Is Essential for Metabolic Research: C57BL/6 and the International Knockout Mouse Consortium. Diabetes 2016;65(1):25–33.

22. [22] Gong WY, Abdelhamid RE, Carvalho CS, Sluka KA. Resident Macrophages in Muscle Contribute to Development of Hyperalgesia in a Mouse Model of Noninflammatory Muscle Pain. J Pain 2016;17(10):1081–1094.

23. [23] Gregus AM, Buczynski MW, Dumlao DS, Norris PC, Rai G, Simeonov A, Maloney DJ, Jadhav A, Holman TR, Xu Q, Wei SC, Fitzsimmons BL, Dennis EA, Yaksh TL. Inhibition of spinal 15-LOX-1 attenuates TLR4-dependent, NSAID-unresponsive hyperalgesia. under review 2018.

24. [24] Gregus AM, Doolen S, Dumlao DS, Buczynski MW, Takasusuki T, Fitzsimmons BL, Hua XY, Taylor BK, Dennis EA, Yaksh TL. Spinal 12-lipoxygenase-derived hepoxilin A3 contributes to inflammatory hyperalgesia via activation of TRPV1 and TRPA1 receptors. Proc Natl Acad Sci U S A 2012;109(17):6721–6726.

25. [25] Gregus AM, Dumlao DS, Wei SC, Norris PC, Catella LC, Meyerstein FG, Buczynski MW, Steinauer JJ, Fitzsimmons BL, Yaksh TL, Dennis EA. Systematic analysis of rat 12/15-lipoxygenase enzymes reveals critical role for spinal eLOX3 hepoxilin synthase activity in inflammatory hyperalgesia. FASEB J 2013;27(5):1939–1949.

26. [26] Gregus AM, Levine IS, Eddinger KA, Yaksh TL, Buczynski MW. Sex differences in neuroimmune and glial mechanisms of pain. Pain 2021;162(8):2186–2200.

27. [27] Haba R, Shintani N, Onaka Y, Wang H, Takenaga R, Hayata A, Baba A, Hashimoto H. Lipopolysaccharide affects exploratory behaviors toward novel objects by impairing cognition and/or motivation in mice: Possible role of activation of the central amygdala. Behav Brain Res 2012;228(2):423–431.

28. [28] Hosseini M, Bardaghi Z, Askarpour H, Jafari MM, Golkar A, Shirzad S, Rajabian A, Salmani H. Minocycline mitigated enduring neurological consequences in the mice model of sepsis. Behav Brain Res 2024;461:114856.

29. [29] Hu YH, Zhang Y, Jiang LQ, Wang S, Lei CQ, Sun MS, Shu HB, Liu Y. WDFY1 mediates TLR3/4 signaling by recruiting TRIF. EMBO Rep 2015;16(4):447–455.

30. [30] Hylden JL, Wilcox GL. Intrathecal morphine in mice: a new technique. Eur J Pharmacol 1980;67(2-3):313–316.

31. [31] Ji RR, Chamessian A, Zhang YQ. Pain regulation by non-neuronal cells and inflammation. Science 2016;354(6312):572–577.

32. [32] Johnson A, Rainville JR, Rivero-Ballon GN, Dhimitri K, Hodes GE. Testing the Limits of Sex Differences Using Variable Stress. Neuroscience 2021;454:72–84.

33. [33] Joshi YB, Di Meco A, Pratico D. Overexpression of 12/15-lipoxygenase increases anxiety behavior in female mice. Neurobiol Aging 2014;35(5):1032–1036.

34. [34] Kaddai V, Gonzalez T, Keslair F, Gremeaux T, Bonnafous S, Gugenheim J, Tran A, Gual P, Le Marchand-Brustel Y, Cormont M. Rab4b is a small GTPase involved in the control of the glucose transporter GLUT4 localization in adipocyte. PLoS One 2009;4(4):e5257.

35. [35] Kaur S, Chen Y, Shenoy SK. Agonist-activated glucagon receptors are deubiquitinated at early endosomes by two distinct deubiquitinases to facilitate Rab4a-dependent recycling. J Biol Chem 2020;295(49):16630–16642.

36. [36] Kehl LJ, Kovacs KJ, Larson AA. Tolerance develops to the effect of lipopolysaccharides on movement-evoked hyperalgesia when administered chronically by a systemic but not an intrathecal route. Pain 2004;111(1-2):104–115.

37. [37] Lacosta S, Merali Z, Anisman H. Behavioral and neurochemical consequences of lipopolysaccharide in mice: anxiogenic-like effects. Brain Res 1999;818(2):291–303.

38. [38] Lenert ME, Avona A, Garner KM, Barron LR, Burton MD. Sensory Neurons, Neuroimmunity, and Pain Modulation by Sex Hormones. Endocrinology 2021;162(8).

39. [39] Levine JD, Lam D, Taiwo YO, Donatoni P, Goetzl EJ. Hyperalgesic properties of 15-lipoxygenase products of arachidonic acid. Proc Natl Acad Sci U S A 1986;83(14):5331–5334.

40. [40] Li JL, Liang YL, Wang YJ. Knockout of ALOX12 protects against spinal cord injury-mediated nerve injury by inhibition of inflammation and apoptosis. Biochem Biophys Res Commun 2019;516(3):991–998.

41. [41] Luo X, Fitzsimmons B, Mohan A, Zhang L, Terrando N, Kordasiewicz H, Ji RR. Intrathecal administration of antisense oligonucleotide against p38alpha but not p38beta MAP kinase isoform reduces neuropathic and postoperative pain and TLR4-induced pain in male mice. Brain Behav Immun 2018;72:34–44.

42. [42] Matsuo N, Takao K, Nakanishi K, Yamasaki N, Tanda K, Miyakawa T. Behavioral profiles of three C57BL/6 substrains. Front Behav Neurosci 2010;4:29.

43. [43] Ming Z, Sawicki G, Bekar LK. Acute systemic LPS-mediated inflammation induces lasting changes in mouse cortical neuromodulation and behavior. Neurosci Lett 2015;590:96–100.

44. [44] Mogil JS. Qualitative sex differences in pain processing: emerging evidence of a biased literature. Nat Rev Neurosci 2020;21(7):353–365.

45. [45] Montilla-Garcia A, Tejada MA, Perazzoli G, Entrena JM, Portillo-Salido E, Fernandez-Segura E, Canizares FJ, Cobos EJ. Grip strength in mice with joint inflammation: A rheumatology function test sensitive to pain and analgesia. Neuropharmacology 2017;125:231–242.

46. [46] Nair DG, Funk CD. A cell-based assay for screening lipoxygenase inhibitors. Prostaglandins Other Lipid Mediat 2009;90(3-4):98–104.

47. [47] Nemoto S, Kubota T, Ohno H. Metabolic differences and differentially expressed genes between C57BL/6J and C57BL/6N mice substrains. PLoS One 2022;17(12):e0271651.

48. [48] Oertel BG, Lotsch J. Clinical pharmacology of analgesics assessed with human experimental pain models: bridging basic and clinical research. Br J Pharmacol 2013;168(3):534–553.

49. [49] Paige C, Barba-Escobedo PA, Mecklenburg J, Patil M, Goffin V, Grattan DR, Dussor G, Akopian AN, Price TJ. Neuroendocrine Mechanisms Governing Sex Differences in Hyperalgesic Priming Involve Prolactin Receptor Sensory Neuron Signaling. J Neurosci 2020;40(37):7080–7090.

50. [50] Papisov MI, Belov VV, Gannon KS. Physiology of the intrathecal bolus: the leptomeningeal route for macromolecule and particle delivery to CNS. Mol Pharm 2013;10(5):1522–1532.

51. [51] Parada CA, Reichling DB, Levine JD. Chronic hyperalgesic priming in the rat involves a novel interaction between cAMP and PKCepsilon second messenger pathways. Pain 2005;113(1-2):185–190.

52. [52] Roosterman D, Cottrell GS, Schmidlin F, Steinhoff M, Bunnett NW. Recycling and resensitization of the neurokinin 1 receptor. Influence of agonist concentration and Rab GTPases. J Biol Chem 2004;279(29):30670–30679.

53. [53] Rudjito R, Agalave NM, Farinotti AB, Lundback P, Szabo-Pardi TA, Price TJ, Harris HE, Burton MD, Svensson CI. Sex- and cell-dependent contribution of peripheral high mobility group box 1 and TLR4 in arthritis-induced pain. Pain 2021;162(2):459–470.

54. [54] Saito O, Svensson CI, Buczynski MW, Wegner K, Hua XY, Codeluppi S, Schaloske RH, Deems RA, Dennis EA, Yaksh TL. Spinal glial TLR4-mediated nociception and production of prostaglandin E(2) and TNF. Br J Pharmacol 2010;160(7):1754–1764.

55. [55] Scholz J, Woolf CJ. The neuropathic pain triad: neurons, immune cells and glia. Nat Neurosci 2007;10(11):1361–1368.

56. [56] Shentu Y, Tian Q, Yang J, Liu X, Han Y, Yang D, Zhang N, Fan X, Wang P, Ma J, Chen R, Li D, Liu S, Wang Y, Mao S, Gong Y, Du C, Fan J. Upregulation of KDM6B contributes to lipopolysaccharide-induced anxiety-like behavior via modulation of VGLL4 in mice. Behav Brain Res 2021;408:113305.

57. [57] Shin J, Cho H, Hwang SW, Jung J, Shin CY, Lee SY, Kim SH, Lee MG, Choi YH, Kim J, Haber NA, Reichling DB, Khasar S, Levine JD, Oh U. Bradykinin-12-lipoxygenase-VR1 signaling pathway for inflammatory hyperalgesia. Proc Natl Acad Sci U S A 2002;99(15):10150–10155.

58. [58] Simon MM, Greenaway S, White JK, Fuchs H, Gailus-Durner V, Wells S, Sorg T, Wong K, Bedu E, Cartwright EJ, Dacquin R, Djebali S, Estabel J, Graw J, Ingham NJ, Jackson IJ, Lengeling A, Mandillo S, Marvel J, Meziane H, Preitner F, Puk O, Roux M, Adams DJ, Atkins S, Ayadi A, Becker L, Blake A, Brooker D, Cater H, Champy MF, Combe R, Danecek P, di Fenza A, Gates H, Gerdin AK, Golini E, Hancock JM, Hans W, Holter SM, Hough T, Jurdic P, Keane TM, Morgan H, Muller W, Neff F, Nicholson G, Pasche B, Roberson LA, Rozman J, Sanderson M, Santos L, Selloum M, Shannon C, Southwell A, Tocchini-Valentini GP, Vancollie VE, Westerberg H, Wurst W, Zi M, Yalcin B, Ramirez-Solis R, Steel KP, Mallon AM, de Angelis MH, Herault Y, Brown SD. A comparative phenotypic and genomic analysis of C57BL/6J and C57BL/6N mouse strains. Genome Biol 2013;14(7):R82.

59. [59] Sorge RE, LaCroix-Fralish ML, Tuttle AH, Sotocinal SG, Austin JS, Ritchie J, Chanda ML, Graham AC, Topham L, Beggs S, Salter MW, Mogil JS. Spinal cord Toll-like receptor 4 mediates inflammatory and neuropathic hypersensitivity in male but not female mice. J Neurosci 2011;31(43):15450–15454.

60. [60] Sorge RE, Mapplebeck JC, Rosen S, Beggs S, Taves S, Alexander JK, Martin LJ, Austin JS, Sotocinal SG, Chen D, Yang M, Shi XQ, Huang H, Pillon NJ, Bilan PJ, Tu Y, Klip A, Ji RR, Zhang J, Salter MW, Mogil JS. Different immune cells mediate mechanical pain hypersensitivity in male and female mice. Nat Neurosci 2015;18(8):1081–1083.

61. [61] Stokes JA, Corr M, Yaksh TL. Spinal toll-like receptor signaling and nociceptive processing: regulatory balance between TIRAP and TRIF cascades mediated by TNF and IFNbeta. Pain 2013;154(5):733–742.

62. [62] Swiergiel AH, Dunn AJ. Effects of interleukin-1beta and lipopolysaccharide on behavior of mice in the elevated plus-maze and open field tests. Pharmacol Biochem Behav 2007;86(4):651–659.

63. [63] Szabo-Pardi TA, Barron LR, Lenert ME, Burton MD. Sensory Neuron TLR4 mediates the development of nerve-injury induced mechanical hypersensitivity in female mice. Brain Behav Immun 2021;97:42–60.

64. [64] Tierney JA, Uong CD, Lenert ME, Williams M, Burton MD. High-fat diet causes mechanical allodynia in the absence of injury or diabetic pathology. Sci Rep 2022;12(1):14840.

65. [65] Trang T, McNaull B, Quirion R, Jhamandas K. Involvement of spinal lipoxygenase metabolites in hyperalgesia and opioid tolerance. Eur J Pharmacol 2004;491(1):21–30.

66. [66] Turnbach ME, Spraggins DS, Randich A. Spinal administration of prostaglandin E(2) or prostaglandin F(2alpha) primarily produces mechanical hyperalgesia that is mediated by nociceptive specific spinal dorsal horn neurons. Pain 2002;97(1-2):33–45.

67. [67] Ulker E, Caillaud M, Patel T, White A, Rashid D, Alqasem M, Lichtman AH, Bryant CD, Damaj MI. C57BL/6 substrain differences in formalin-induced pain-like behavioral responses. Behav Brain Res 2020;390:112698.

68. [68] Witkin LR, Zylberger D, Mehta N, Hindenlang M, Johnson C, Kean J, Horn SD, Inturrisi CE. Patient-Reported Outcomes and Opioid Use in Outpatients With Chronic Pain. J Pain 2017;18(5):583–596.

69. [69] Woller SA, Corr M, Yaksh TL. Differences in cisplatin-induced mechanical allodynia in male and female mice. Eur J Pain 2015;19(10):1476–1485.

70. [70] Woller SA, Ravula SB, Tucci FC, Beaton G, Corr M, Isseroff RR, Soulika AM, Chigbrow M, Eddinger KA, Yaksh TL. Systemic TAK-242 prevents intrathecal LPS evoked hyperalgesia in male, but not female mice and prevents delayed allodynia following intraplantar formalin in both male and female mice: The role of TLR4 in the evolution of a persistent pain state. Brain Behav Immun 2016;56:271–280.

71. [71] Yaksh TL, Woller SA, Ramachandran R, Sorkin LS. The search for novel analgesics: targets and mechanisms. F1000Prime Rep 2015;7:56.

72. [72] Yekkirala AS, Roberson DP, Bean BP, Woolf CJ. Breaking barriers to novel analgesic drug development. Nat Rev Drug Discov 2017;16(8):545–564.

